# Molecular Sequestration Enhances Precision in Timing of Intracellular Events

**DOI:** 10.1101/2024.05.23.595626

**Authors:** Kuheli Biswas, Supravat Dey, Abhyudai Singh

## Abstract

Expressed gene products often interact ubiquitously with binding sites at nucleic acids and macromolecular complexes, known as decoys. Binding of transcription factors (TFs) to decoys is shown to be an indirect but crucial way to control the dynamics and stochasticity in gene regulation. Here, we explore how such decoys impact the timing of intracellular events, as captured by the time taken for the levels of a given TF to reach a critical threshold level, known as *first passage time* (FPT). As binding introduces nonlinearity, an exact mathematical analysis is challenging. However, assuming quasi-static equilibrium (QSE) for binding/unbinding reactions, we can analytically discern the impact of decoys on FPT statistics by using small noise approximation (SNA) and reformulating the FPT question in terms of a suitable variable whose dynamics is linear. The stability of the decoy-bound TFs influences the impact of decoys on FPT statistics. The presence of decoys makes the mean FPT long and even longer for unstable bound TFs. Decoys reduce noise in FPT, and stable decoy-bound TFs offer greater timing precision with less expression cost than their unstable counterparts. Interestingly, when both bound and free TFs decay at the same rate, the noise in FPT is not directly influenced by the number of decoys or their binding affinities. We verify these results by performing exact stochastic simulations. These results have important implications for the precise temporal scheduling of events involved in the functioning of biomolecular clocks, development processes, cell-cycle control, and cell-size homeostasis.

## I. INTRODUCTION

Sequestration of gene products at genomic sites and in phase-separated particles critically shapes the stochastic dynamics of biomolecular circuits defining cellular responses to diverse stimuli (1–6). For example, Transcription Factors (TF) not only bind to their target promoters but also bind promiscuously to “decoy sites” scattered across the genome (7–9). When binding to such decoys stabilizes an otherwise actively degraded TF, then decoys have been shown to buffer stochastic variation in the levels of free (unbound) TF (10–12). In contrast, when binding does not impact TF stability, decoys can function as noise amplifiers (13). This latter scenario can occur for stable TFs whose concentrations are diluted from cellular growth.

While both experimental and computational work have explored the effects of sequestration mechanisms on stochastic variation in gene product levels, their impact on event timing remains mostly unexplored. Some theoretical and computational works have shown that the sequestration of TFs with decoys plays diverse roles in controlling the mean and stochastic dynamics in various genetic circuits such as oscillators (14–19) and toggle switches (19). Recent studies have shown that decoy binding can crucially affect an event’s mean first passage time (FPT) in auto-regulatory circuits (20), and play an important role in precision control of bacterial replication initiation (21). In this contribution, we explore the influence of decoys on the timing noise of events, employing FPT formalism. This analysis considers events triggered by the accumulation of a specific protein to a predefined threshold level. Several works have derived analytical expressions for FPT statistics for different models of stochastic gene expression, including models with feedforward and feedback regulation (22–25), post-transcriptional regulation (26, 27), miRNA-mediated regulation of protein translation (28), providing rich insight into event timing in the context of different biological phenomena such as cell-cycle regulation (29), lysis timing of phages (30–32), development (33), cell-state transitions (34, 35), and neuronal firing of action potentials (36, 37). Here we systematically explore how decoy-based sequestration of gene products impacts stochasticity in FPT using well-established analytical approximations corroborated with exact stochastic simulations.

Understanding the variability in the timing of gene expression is crucial for unraveling fundamental biological mechanisms at the single-cell level. Identifying the regulatory strategies that can modulate this variability is essential for decoding the underlying design principles of cellular regulatory networks that govern timing. Advancements in experimental techniques, notably fluorescence time-lapse microscopy, often integrated with microfluidic platforms to maintain cells in a precisely controlled environment across multiple generations, are poised to significantly enhance data accessibility on gene expression timing and its variabilities. Consequently, a robust theoretical framework elucidating the dynamics of timing fluctuations is imperative. Such a framework is necessary to effectively interpret forthcoming data, guide the development of targeted experimental designs, and facilitate the engineering of synthetic biological circuits with tailored timing characteristics.

## II. MODEL FORMULATION

To analytically explore the impact of decoy binding on the FPT statistics we consider a simplified gene expression model for the TF protein, excluding the dynamics of mRNA and gene activity. The schematic of the model for TF synthesis and its interaction with decoy binding sites are shown in Fig. 1. Consistent with empirical observations, the TF synthesis is characterized as a stochastic burst event with a burst frequency denoted as *k* and a burst amplitude (size) represented by *B* (38–47). We consider a general form for the burst size distribution throughout the paper, except for plotting and simulation purposes where we consider a geometrical distribution grounded in prior research (48–51). A TF molecule then decays with rate *γ*_*f*_.

**FIG. 1.**
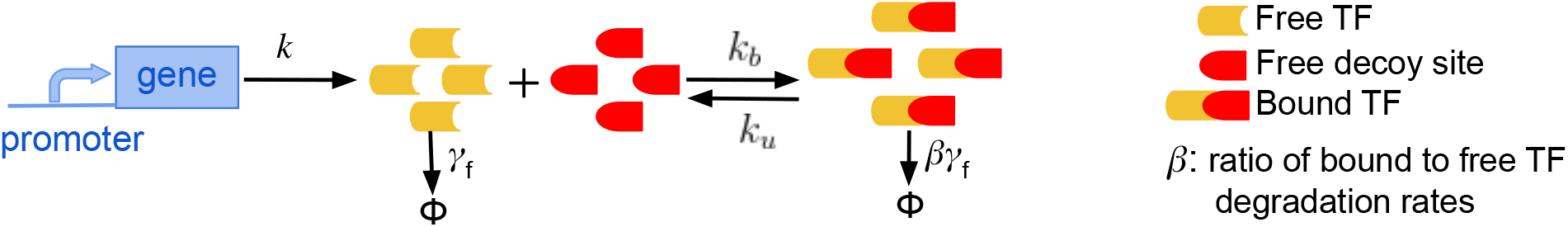
Schematic of TF synthesis model in the presence of decoy binding sites. **(a)** The synthesis of free TFs occurs in stochastic bursts with rate *k*, which reversibly bind/unbind to *N* decoy binding sites with rates *k*_*b*_ and *k*_*u*_. Both the free and bound TFs are subject to degradation with rates *γ*_*f*_ and *βγ*_*f*_, respectively.

An expressed TF molecule binds reversibly to a decoy from a pool of total *N* decoy sites to form a decoy-bound complex. The binding and unbinding rates are *k*_*b*_ and *k*_*u*_, respectively, and *N* is kept fixed in our study. Experimental evidence suggests that the decay of decoy-bound TFs is context-dependent. For some TFs such as p53 (52) and MyoD (53), the decoy binding protects TFs from degradation. While some other TFs decoy-binding promotes degradation such as VP16 in *Saccharomyces cerevisiae* via ubiquitin-mediated proteolysis (54). It has been shown theoretically that the stability of bound TFs plays crucial roles in the deterministic and stochastic dynamics of the nonregulatory (13), auto-regulatory(10), and oscillatory gene circuits (17). Here, we assume the degradation of a TF molecule in the bound state is *βγ*_*f*_. We will focus on two values of *β*=1 and 0, where *β*=0 corresponds to no decay of bound TF, and *β*=1 corresponds to equal degradation rates for both the free and bound TF.

Our TF expression and sequestration model at decoy binding sites is based on the standard stochastic formulation of chemical kinetics (55, 56). The model is comprised of five following events that occur probabilistically at exponentially distributed time intervals

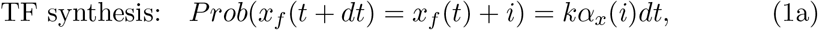

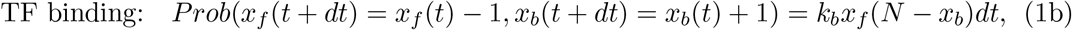

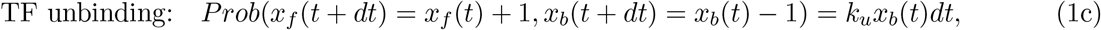

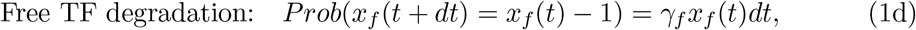

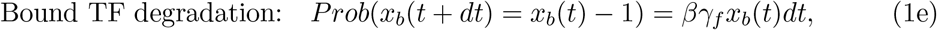

where *α*_*x*_(*i*) is the probability distribution for burst size *B* = *i. x*_*f*_ (*t*), *x*_*b*_(*t*) and *x*(*t*) := *x*_*f*_ (*t*)+*x*_*b*_(*t*) denote the level of free, bound and total (free + bound) TF at time *t* inside the cell, respectively. To study the role of decoy binding sites on the statistics of event timing, our goal is to characterize the dynamical moments of free TF numbers as a function of the number of decoy sites *N*. The first and second moments of a stochastic variable *Y* at time *t* are denoted as ⟨*Y* ⟩ and ⟨*Y* ^2^⟩, respectively. The angular bracket ⟨·⟩ represents ensemble averages, which are obtained by averaging over many trajectories in simulations. The noise in *Y* is quantified by its coefficient of variation square at time *t*, which is denoted by 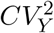. For readers’ convenience, we provide a list of model parameters and their description in Table I.

**TABLE I.** Summary of notation used.

| Symbol | Description | Symbol | Description |
| --- | --- | --- | --- |
| $k$ | TF burst frequency | $B$ | TF Bust size |
| $k_b$ | TF binding rate | $k_u$ | TF unbinding rate |
| $k_d$ | Dissociation constant ($k_u/k_b$) | $\gamma_f$ | Free TF degradation rate |
| $\beta\gamma_f$ | Bound TF degradation rate | $N$ | Total decoy binding sites |
| $x_f$ | Free TF number at time $t$ | $\langle Y \rangle$ | Mean value of variable $Y$ at time $t$ |
| $x_b$ | Bound TF number at time $t$ | $CV_Y$ | Coefficient of variation of $Y$ at time $t$ |
| $x$ | Total TF number ($x_f + x_b$) at time $t$ | $X$ | Threshold number of free TFs |
| $T$ | First passage time | $\mathcal{T}$ | $:= \gamma_f T$ FPT in the dimensionless unit |
| $x_0$ | $:= k\langle B \rangle / \gamma_f$, The stead-state mean of TF numbers for no Decoy, $N = 0$ | | |
| $g$ | $g(X, N, k_d)$, fraction of free TFs in total TFs at threshold crossing | | |

## III. FPT STATISTICS IN THE ABSENCE OF DECOYS

The first passage time (FPT) or event timing in our study is defined as the minimum time required for the free TF level to reach a critical threshold for the first time to trigger some downstream processes. More precisely, starting from zero initial conditions, the FPT is mathematically described by the following random variable

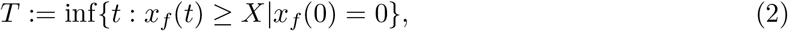

where *X* is the critical threshold level of free TFs needed for some events to occur. FPT is a stochastic variable since the free TF number fluctuates over time. When the fluctuations in the free TF level are small, the mean FPT, ⟨*T* ⟩ is approximately the time required for the mean free TF level to reach the threshold for the first time, i.e.,

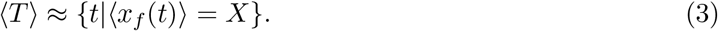

In the absence of decoys, the mean dynamics of free TFs are usually straightforward to obtain from linear biochemical systems given as follows (57)

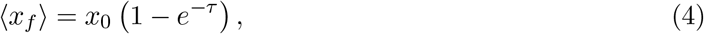

which increases monotonically with time, asymptotically approaching the steady-state value *x*_0_ := *k*⟨*B*⟩*/γ*_*f*_. For simplicity, we use dimensionless time, *τ* = *γ*_*f*_ *t*, implying time measurement in the unit of lifetime of a free TF molecule.

The noise in FPT is quantified by the squared coefficient of variation (*CV* ^2^), which is the ratio of the variance and the mean squared in FPT. By using the small noise approximation (SNA) in free TF level one can compute the FPT noise as follows (see Appendix A)

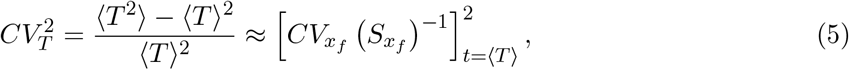

where

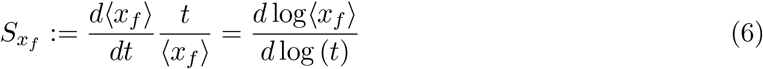

is the dimensionless log sensitivity of the mean free TF level with respect to time. In the absence of decoys, the noise dynamics of free TFs is given as follows (57)

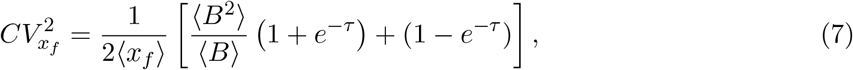

which asymptotically reaches the steady-state value ⟨*B*^2^⟩*/*⟨*B*⟩*x*_0_. We note that Eq. (5) leads to the same mathematical expression as in Ref (58) obtained by a different approach by following the simple geometric argument in TF number dynamics.

Now, using the dynamics of mean and noise of free TFs in the definition of mean FPT (Eq. 3) and FPT noise (Eq. 5) one can obtain

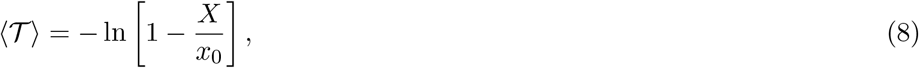

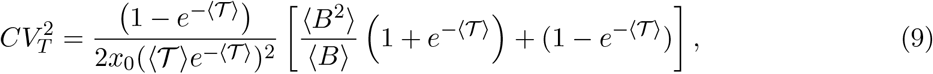

which is the same expression as in the Ref. (58). We use mean FPT in the dimensionless unit ⟨𝒯 ⟩ = *γ*_*f*_ ⟨*T* ⟩. One significant feature emerges from Eq. (9) is that FPT noise exhibits a nonmonotonic behavior with mean FPT and the optimal value of normalized mean FPT is 𝒯 = ln 2 for geometric burst size distribution (58). The identification of the optimal mean FPT at 𝒯 = ln 2 is further substantiated by recent theoretical advances, where the lysis time has been analytically modeled within the FPT framework (59). Their precise analytical calculations, devoid of any approximations, are in remarkable agreement with the experimental measurements of lysis times observed in variants of *λ* phages (31).

## IV. FPT STATISTICS IN THE PRESENCE OF DECOYS

In the presence of decoys, the average dynamics for free and bound TF numbers are nonlinear and will be expressed as follows (12, 13):

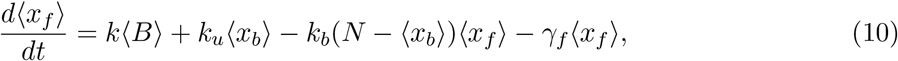

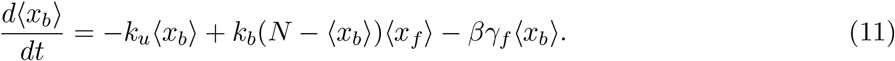

The nonlinearity in the dynamics of free and decoy-bound TFs makes calculating FPT statistics challenging. However, as we will see below, in some limits one can bypass this nonlinearity by considering the dynamics of total TFs, *x* = *x*_*f*_ + *x*_*b*_ as follows

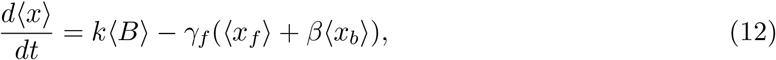

and reformulating the FPT question in terms of total TFs.

We start with the assumption *k*_*u*_*x*_*b*_=*k*_*b*_(*N* − *x*_*b*_)*x*_*f*_, i.e., the binding/unbinding reactions occur at a faster rate compared to TF synthesis and degradation. This assumption, known as the quasi-static equilibrium (QSE) or adiabatic limit, is a prevalent approach in gene expression modeling (60–62), supported by recent experimental evidence (63). Using this approximation we get the relation between total and free TF numbers as follows

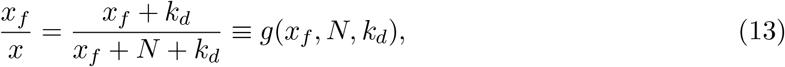

where 0 ≤ *g*(*x*_*f*_, *N, k*_*d*_) ≤ 1 and it signifies the fraction of free TF molecules. For notational convenience, we use *g*(*X, N, k*_*d*_) as *g* henceforth.

Based on Eq. (13) we reformulate the FPT in terms of the dynamics of the total TF level. Instead of locating the time for the free TF level to reach a threshold *X* for the first time, we find the time when the total TF level first time hits a normalized threshold *X/g*. More precisely, the modified FPT is mathematically described by the following random variable

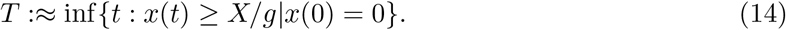

We numerically demonstrate that this assumption works well when the noise in the system is relatively small. In Fig. (2), we have plotted the simulated trajectories and distributions of actual FPT (Eq. 2) and modified FPT (Eq. 14). Not only the mean FPT, but the full distributions of FPT from these two definitions appear to be identical. This approximation even persists when the binding/unbinding reactions are slow, of the order of TF degradation rate (see Appendix Fig. S1). However, in the limit of a small free TF threshold, *X* ≪ *x*_0_ these two distributions deviate from each other, and the approximation is no longer valid (see Appendix Fig. S2).

Following the same SNA approximation in *x* as mentioned before in Eq. (3), we can write the modified mean FPT as

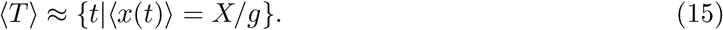

Similar to Eq. (5), the FPT noise will be as follows

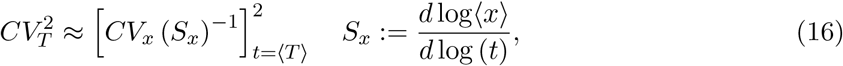

which depends on the noise dynamics of total TFs 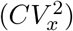 and the dimensionless log sensitivity of the mean total TF level with respect to time (*S*_*x*_) (see Appendix A). So, using SNA and fast binding/unbinding approximations, we can obtain analytical expressions for the mean and noise of free TFs’ FPT from the total TFs’ moment dynamics. Next, we will focus on two key regimes characterized by *β*=0 implying stable decoy-bound TFs and *β*=1 indicating the same degradation rate of free and decoy-bound TFs.

### Case -I: Both bound and free TFs decay at the same rate (β = 1)

Using *β*=1 in Eq. (12) one can obtain the average dynamics of the total TF number, which is the same as the dynamics of average TF number for no decoy (Eq. 4). Now using the modified definition of mean FPT in Eq. (15), one can obtain an expression from mean FPT as follows

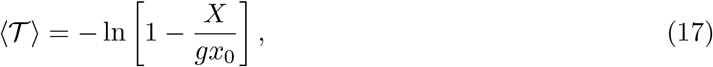

which reduces to Eq. (8) for no decoys as the fraction of free TFs *g*=1 when *N* =0. The mean FPT increases monotonically with *N* and diverges for large *N* as *g* → 0 when *N* → ∞ (Fig. 3a). The analytical form of mean FPT obtained by using the modified threshold on total TFs (Eq. 15) agrees well with the exact stochastic simulation algorithm (SSA) (64) (see Fig. 3a) of the decoy model in Eq. (1) that considers the actual free TF number threshold (Eq. 2).

**FIG. 2.**
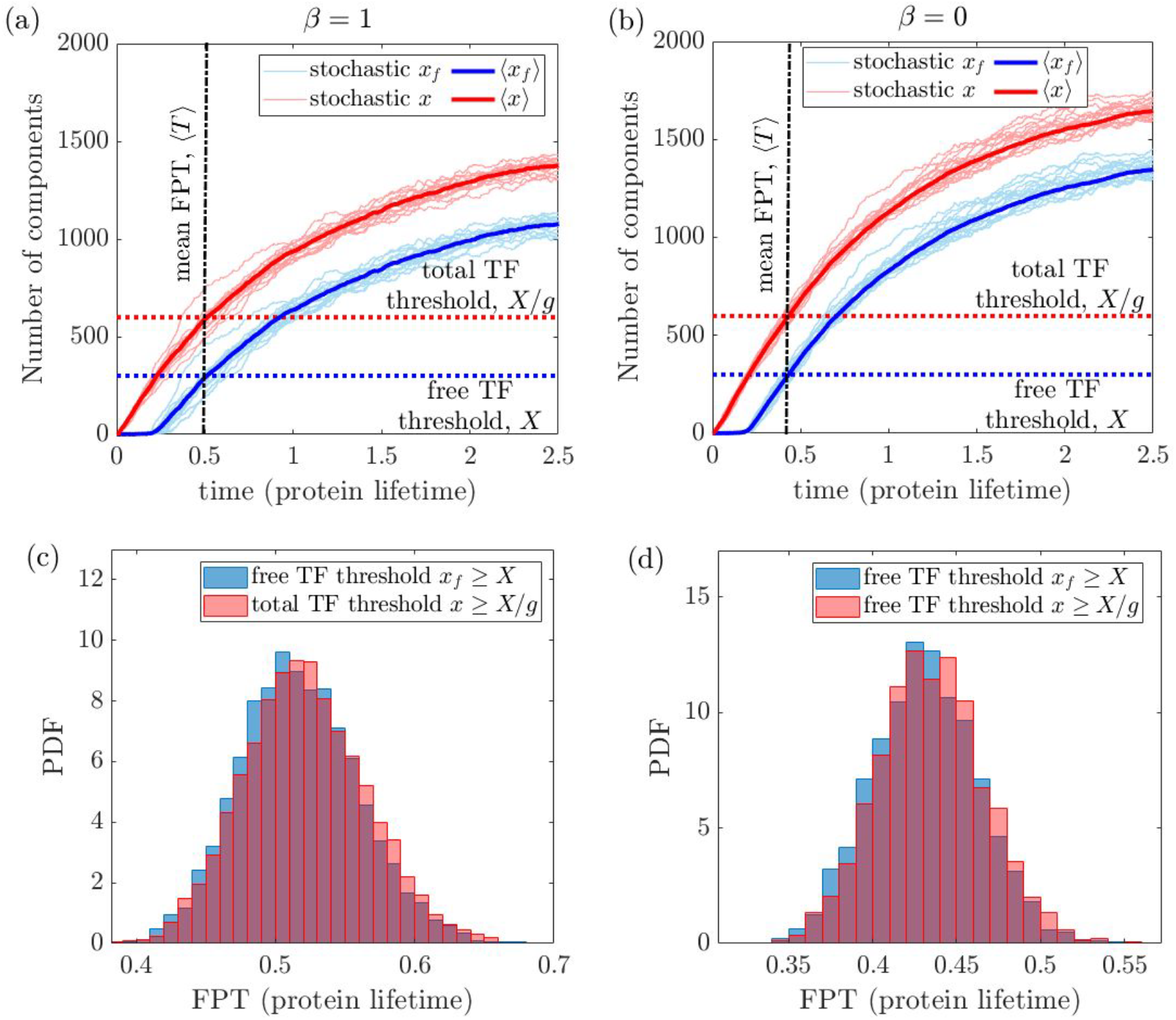
FPT statistics of free TF level reaching a threshold *X* is approximated by FPT statistics of total TF level reaching a normalized threshold *X/g*. Simulated stochastic and mean trajectories of total (*x*) and free (*x*_*f*_) TFs for **(a)** *β* = 1 and **(b)** *β* = 0 along with threshold of *x*_*f*_ is *X* and *x* is *X/g*. Simulated FPT distributions from both thresholds in Eqs. (2) and (14) will show a significant overlap for **(c)** *β* = 1 and **(d)** *β* = 0. We assume the shifted geometric distribution of burst size, i.e., ⟨*B*^2^⟩ = 2⟨*B*⟩^2^ + ⟨*B*⟩ with parameter values: *N* = 300, *X* = 300, *γ*_*f*_ = 1, *k*_*b*_ = *k*_*u*_ = 50, ⟨*B*⟩ = 10, *x*_0_ = 1500. We use the stochastic simulation algorithm (SSA) (64) over 5 × 10^3^ realizations on model Eq. (1).

**FIG. 3.**
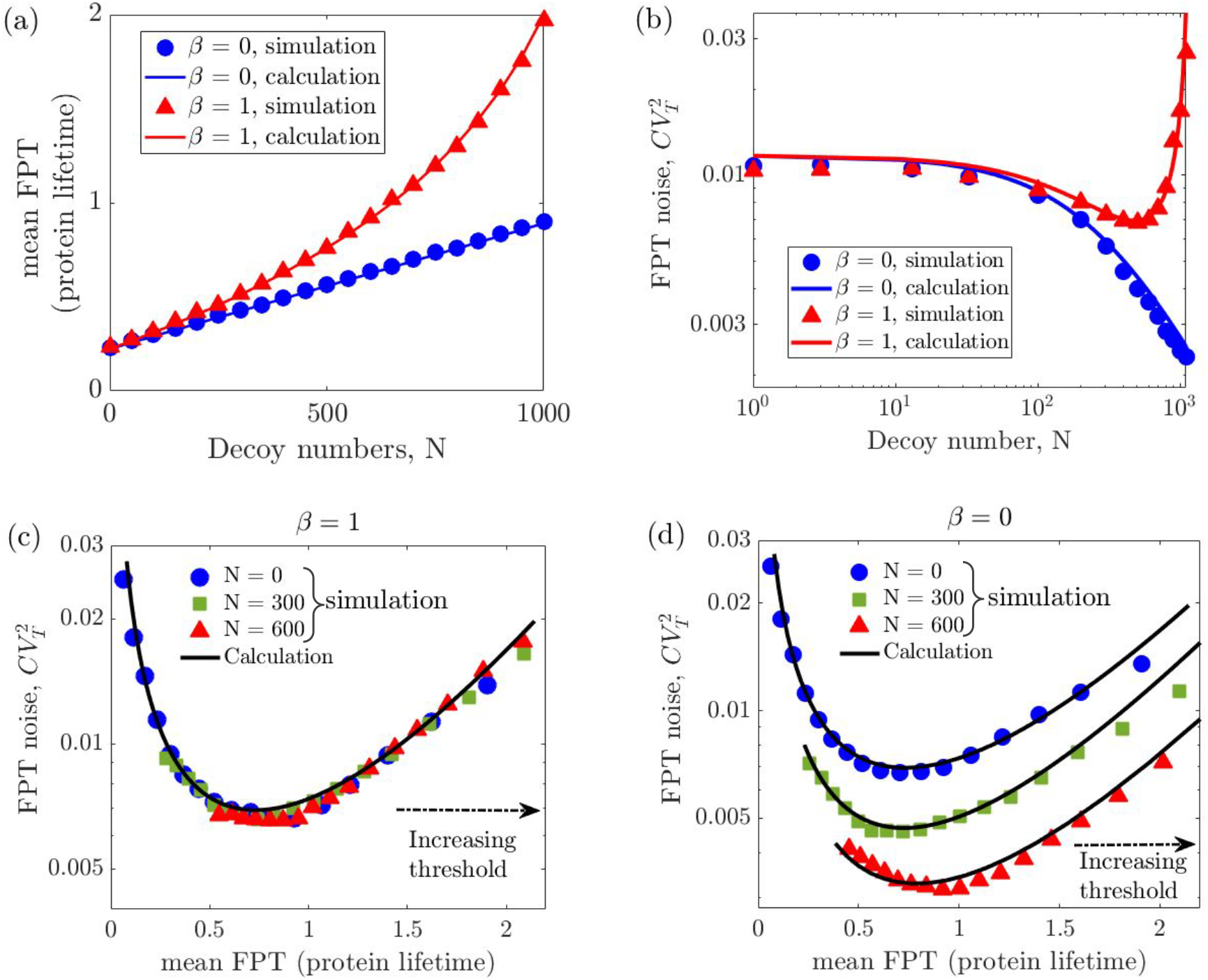
Stability of bound TFs distinctly influences the FPT statistics. Variation of **(a)** mean FPT and **(b)** FPT noise with decoy number for *β* =1 Eqs. (17), (18) (red triangles) and *β* =0 Eqs. (20), (22) at fixed threshold value *X* = 300. Variation of FPT noise with mean FPT for **(c)** *β* = 1 and **(d)** *β* = 0 for three different values of *N*; *N* = 0, 300, and 1000. For *β* = 1, the relation between ⟨*T* ⟩ and 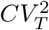 follows a master curve Eq. 18 for all *N* values, which deviates for *β* = 0, for nonzero *N* values (Eq. 22). For the simulation, we use the stochastic simulation algorithm (SSA) (64) over 5 × 10^3^ realizations using Eqs. (1), and (2). For each *N*, we change the value of ⟨*T* ⟩ by changing the threshold *X*. We assume the shifted geometric distribution of burst size, i.e., ⟨*B*^2^⟩ = 2⟨*B*⟩^2^ + ⟨*B*⟩ with parameter values: *γ*_*f*_ = 1, *k*_*b*_ = *k*_*u*_ = 50, ⟨*B*⟩ = 10, *x*_0_ = 1500.

Now, for the analysis of the FPT noise of free TF, we need the noise dynamics of the total TF number as mentioned in Eq. (16). In the limit of *β*=1, the noise in total TF numbers is the same as no decoy case, which is given in Eq. (7) (see Appendix B). Using the expressions of mean and noise dynamics of total TF numbers in Eq. (16) one can obtain the mathematical form of FPT noise as follows

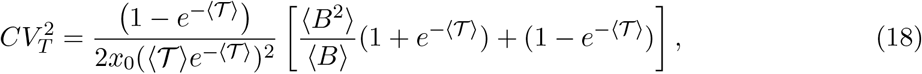

which is the same as Eq. (9) for no decoy case. In Eq. (18),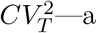 measure of FPT noise—implicitly depends on *N* via ⟨𝒯 ⟩. This implies that the relation between the FPT mean and noise remains invariant of *N* when *β*=1 as presented in Fig. 3c. So, for *β*=1, with different *N* values, ⟨*T* ⟩ and hence 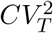 will change in such a way that it will follow a master curve, which is Eq. (18). It is remarkable to note that despite the influence of decoy binding on the dynamics of free TFs, it does not modify the relationship between the FPT mean and noise. The optimal value of mean FPT is ⟨𝒯⟩ = ln 2, similar to the no decoy case as mentioned before in sec. (III). The approximated FPT noise from our analysis agrees well with the exact SSA (64). Increasing the number of decoys results in a higher mean FPT (Fig. 3a) and thus impacts the FPT noise according to Eq. (18). The dependence of FPT noise on the number of decoys is depicted in Fig. (3)b.

### Case -II: Bound TFs are protected from degradation (β = 0)

Using *β*=0 in Eq. (12) one cannot exactly solve the differential equation for ⟨*x*⟩. However, in the limit of *k*_*d*_ ≪ *x*_*f*_, i.e., when all decoy sites are bound with TFs (*x*_*b*_ ≈ *N*) one can get an approximated dynamics of the total TF number as follows

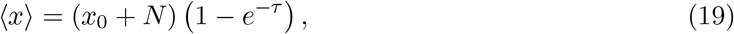

which is the same as the dynamics of the free average TF number for *N* =0 (Eq. 4). Now using the modified definition of FPT in Eq. (15), one can obtain an expression from mean FPT as follows

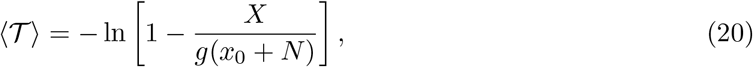

which reduces to Eq. (8) for no decoys. For *β*=0, though the mean FPT increases with *N*, it does not show a diverging trend like *β*=1 (Eq. 17) as *g*(*x*_0_ + *N*) → (*X* + *k*_*d*_) when *N* → ∞. Our approximated calculation for mean FPT in Eq. (20) agrees well with the stochastic simulation algorithm (SSA) (64) of the decoy model in Eq. (1) (Fig. 3a).

Now, for the analysis of noise in free TF numbers’ FPT, we need the noise dynamics of the total TF number as mentioned before in Eq. (16). In the limit of *β*=0, the noise in total TF numbers is given as follows (see Appendix B)

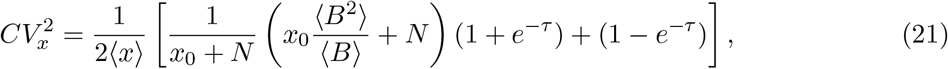

which converges to Eq. (7) for *N* =0. Using the expressions of ⟨*x*⟩ (Eq. 19) and *CV*_*x*_ (Eq. 21) in Eq. (16) one can obtain the mathematical form of FPT noise as follows

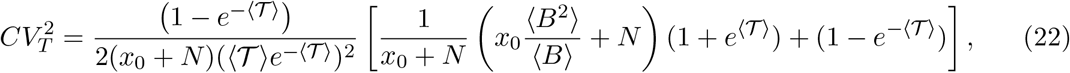

which is an explicit function of *N*, unlike for the case of *β*=1 in Eq. (18). The variation of FPT noise with decoys is shown in Fig. 3b. Eq. (22) demonstrates that the relationship between the FPT mean and noise deviates from the master curve observed in the absence of decoys as 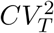 is inversely proportional to the number of decoys (*N*). This analysis is also supported by the stochastic simulation algorithm (SSA) results shown in Fig. (3)d.

## V. FPT NOISE FOR A FIXED MEAN FPT

In the previous section, we observe how the decoys alter both the mean FPT and the noise for a given production rate, decay rate, and binding affinity of transcription factors. The response of living cells to the presence of decoys influencing event timing remains unclear. It can be hypothesized that cells may maintain a constant mean FPT. In this section, we explore the behavior of FPT noise while holding the mean FPT constant. Consequently, to keep the mean FPT constant while varying *N*, it becomes necessary to modify certain parameters as ⟨T ⟩ = *f* (*N, X, x*_0_, *k*_*d*_) (Eqs. 17, 20 for *β*=1 and 0). Here, we focus on adjusting burst frequency (*k*) hence *x*_0_(= *k*⟨*B*⟩) and threshold (*X*). However, increasing burst frequency results in a higher load of TFs in the cell. Similarly, raising the threshold level leads cells to produce more TFs. Therefore, enhancing both the burst frequency and the threshold level increases the TF load on the cell, which we describe qualitatively as being more “expression cost” for the cell. Moving forward, we will explore how variations in burst frequency and threshold influence the timing precision of cellular processes, particularly in scenarios involving the presence of decoys, for both *β*=1 and 0.

First, we will focus on tuning the threshold, *X* to maintain a fixed mean FPT. For both *β* values, the threshold decreases linearly with increasing decoys (Fig. 4b). A reduced threshold requirement means fewer TFs need to be produced, implying lesser expression cost for cells. Moreover, when *β*=0, it can be seen from Eq. 22 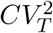 monotonically decays with *N* and asymptotically follows a 1*/N* relationship (Fig. 4a). Therefore, for stable decoy-bound TFs (*β*=0), decoy binding enhances both precision and cost-effectiveness for cells, provided the threshold is tuned to keep the mean FPT constant (see Fig.4b). In contrast, for *β*=1, the noise in FPT is maintained at the same level observed without decoys, regardless of the increase in *N* as seen in Fig. 4a. This outcome occurs as 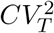 in Eq. (18) implicitly depends on *X* and *N* through ⟨*T* ⟩, which is kept fixed.

**FIG. 4.**
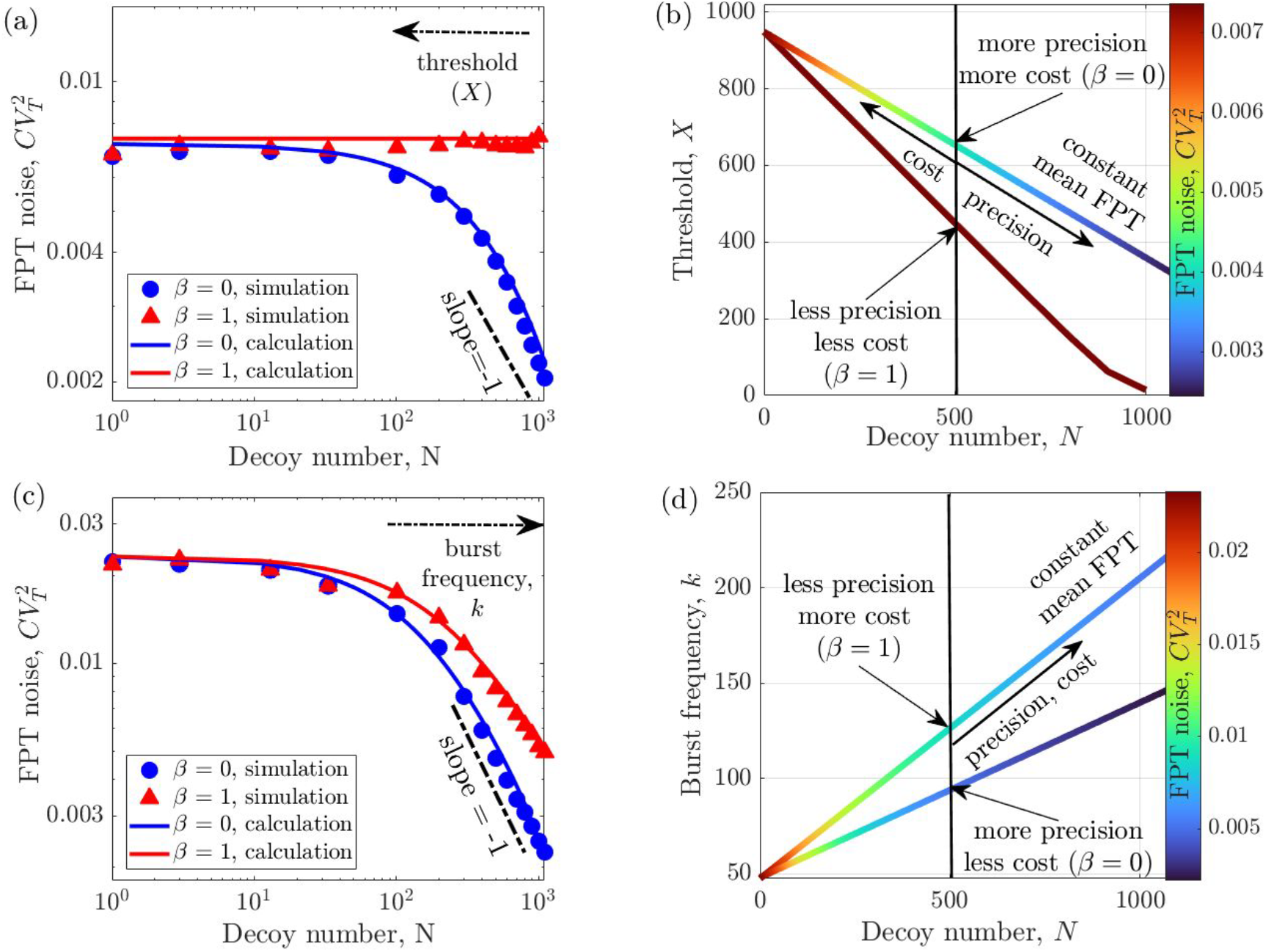
Decoys enhance timing precision at the expense of higher expression cost for a fixed mean FPT. Variation of the FPT noise with decoy number for fixed ⟨T ⟩ = 1 by changing **(a)** threshold for a fixed burst frequency *k* = 150 and varying **(c)** burst frequency for a constant threshold *X* = 300. For **(a)** and **(c)** we tune, respectively, *X* and *k* by using Eq. (17) for *β* = 1 and Eq. (20) for *β* = 0. The parameters used for figures: *γ*_*f*_ = 1, *k*_*b*_ = *k*_*u*_ = 50, ⟨*B*⟩ = 10. We use the SSA (64) over 5 × 10^3^ realizations for simulation. Contours representing a constant mean FPT of 1 in the **(b)** *N* − *X* and **(d)** *N* − *k* plane are depicted for *β* = 1 and *β* = 0. The contours are color-coded by 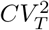 quantifying FPT noise levels. Precision increases in the direction of the arrow, indicating lesser noise. On the same contour, higher thresholds and burst frequency increase “expression costs” due to elevated TF loads. While not quantified, the expression cost qualitatively increases in the direction of the arrow.

Next, we will focus on tuning burst frequency to maintain a constant mean FPT fixed for varying values of *N*. Using the mean FPT equations for *β*=1 (Eq. 17) and *β*=0 (Eq. 20), we can precisely determine *x*_0_ and, subsequently, the burst frequency, *k*=*x*_0_*/*⟨*B*⟩ for a fixed m ean FPT while *N* is changing. For both scenarios, the burst frequency increases linearly with *N*, as shown in Fig. 4(d). This increase in burst frequency increases transcription cost (65) hence expression cost. Moreover, linearly increasing burst frequency with decoy numbers reduces FPT noise as 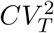 inversely correlates with burst frequency both for *β*=1 (Eq. 18) and *β*=0 (Eq. 22). Hence FPT noise decays as 1*/N* when we tune burst frequency to maintain a fixed mean FPT. The exact analytical forms of 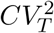 for both *β* values are plotted in Fig. 4(c), which have −1 slope in the log-log axes (black dashed line for the eye guide). Our approximated analytical calculation matches the exact stochastic simulations (64) of the model described in Eqs. (1) and (2). Therefore, decoy binding enhances timing precision at the expense of higher expression cost (see Fig.4d). However, for a specific number of decoy binding sites (*N*), stable bound TFs (*β*=0) offer greater timing precision with lower expression costs compared to unstable bound TFs (*β*=1) as detailed in Fig. 4d.

## VI. DISCUSSION

In this study, we have systematically investigated how sequestration of TFs by decoy sites impacts the statistics of event timing, a scenario not extensively explored in the existing literature. Specifically, we have studied the first passage time (FPT) statistics, the time required for free TF numbers to cross a certain threshold value. We have obtained approximated analytical expressions for the mean and noise of FPT. The noise is quantified by the square of the coefficient of variation 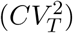 in FPT. As the decoy binding introduces nonlinearity, the analytical tractability of the problem usually becomes challenging, especially for the dynamics not only for the mean but also for the noise. Our analytical calculations of FPT moments are based on two key approximations: (1) quasi-static equilibrium (QSE), i.e., fast binding/unbinding reactions, and (2) small noise approximation (SNA) in TF levels. We have shown that we can eliminate nonlinearity by looking at the dynamics of the total TF level and reframing the FPT question in terms of the total TF level reaching an equivalent threshold. Our approximated simple analytical results provide better insights for understanding FPT noise and agree with exact stochastic simulations. Without decoys, the exact analytical calculations for FPT statistics are possible (31). However, the complexity in the mathematical expression for exact formulas may prohibit simple intuitive insights that one achieves via SNA approximation (58). While this contribution focuses on TFs binding to genomic decoy sites, this framework applies to other classes of proteins, for example, RNA-binding proteins binding to sites on RNA (66–68).

Our investigation indicates that the stability of decoy-bound TFs is critical in determining FPT statistics. Earlier studies have shown that stable bound TFs decrease gene expression variability at steady-state, whereas their degradation increases fluctuations (10, 13). In the context of biomolecular clocks, however, the degradation of bound TFs can enhance oscillatory behavior and reduce noise, while protecting these TFs from degradation can disrupt sustained oscillations in gene expression (17). This study shows that the mean FPT increases with the number of decoy binding sites and this increment is rapid if bound TFs are unstable (Fig. 3a). Importantly, when the degradation rates for free and bound TFs are equal (*β*=1), FPT noise does not explicitly depend on the number of decoys but is influenced indirectly via the mean FPT. In this scenario, adjusting burst frequency (threshold) to maintain a constant mean FPT effectively reduces FPT noise (regulates) with decoys compared to the no-decoy scenario (see Figs. 4a, c). In contrast, for stable bound TFs (*β*=0), decoys consistently reduce FPT noise, as it directly correlates with the inverse function of the number of decoy sites (Eq. 22). On the other hand, while increasing decoys enhances the precision of event timing, it comes at the cost of a higher total protein load and consequently, increased expression cost.

The analytical formulas for the noise give theoretical insights into the role of decoys on the noise of FPT. These are derived by assuming small copy number fluctuations around the statistical mean and then taking fast bind/binding limit. Using numerical exact stochastic simulations, we have shown that the main results agree with the theory. The quantitative match between theory and simulations can be poor where the fluctuations are large. We numerically have found that results are not very sensitive to fast binding/unbinding limits (see Appendix Fig. S3). The objective of obtaining more accurate theoretical results is a matter of future work. Our novel finding of the dual role of decoys as noise regulators/buffers in event timing encourages more investigation into the regulatory function of decoys in complex gene networks. In our prior research, we used bacteriophage *λ* as a model system for studying event timing in individual cells. Here, an easily observable event (cell lysis) is the result of the expression and accumulation of a single protein (holin) in the *E. coli* cell membrane up to a threshold level (30, 32, 59), and precision in event timing is tied an optimal timing of phage-mediated cell lysis (69). The lysis pathway also consists of another protein, antiholin that binds to holin and prevents holin from participating in hole formation (70, 71). Expressing antiholin in titratable amounts from a plasmid could be an interesting setup to experimentally investigate the role of decoys on intracellular event timing and connect these results with the theoretical foundation setup here. Finally, it will also be interesting to consider different forms of parametric fluctuations coupled with the intrinsically noisy expression, for example, the threshold level itself could be random in single cells (72).

## Appendix A: Relation between CVs of two stochastic variable

If we have two stochastic variables 𝒳 and 𝒴 and we know the relation between them as

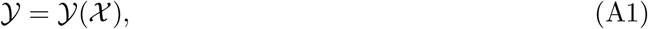

then using the small noise approximation (SNA) in 𝒳 around the mean value ⟨𝒳⟩ we can write

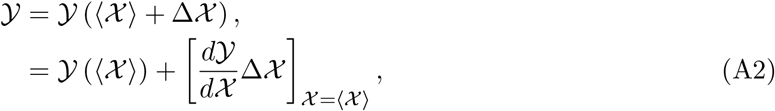

where Δ𝒳 is the white noise in stochastic variable 𝒳 with mean, ⟨Δ𝒳⟩ = 0 and variance,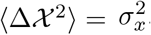. Using this one can calculate the mean and variation of the stochastic variable 𝒴, respectively, as follows

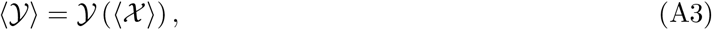

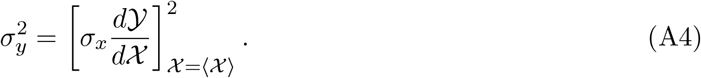

So, the coefficient of variation of the stochastic variable 𝒴 will be

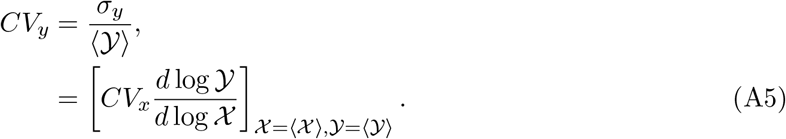

We define

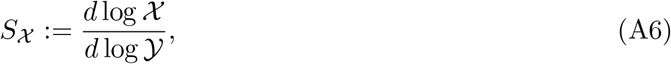

which is the dimensionless log sensitivity of stochastic variable 𝒳 with respect to 𝒴 evaluated at mean values. *CV*_*x*_ is the coefficient of variation of the stochastic variable 𝒳, which is *σ*_*x*_*/*⟨𝒳⟩. This equation is the same as Eq. (5) with 𝒴 ≡ *t*, FPT and 𝒳 ≡ *x*_*f*_, free TF number.

## Appendix B: CV dynamics of total TFs for both *β* values

When both free TF and bound TF decay at the same rate (*β* = 1), the dynamics of the total TF is given by,

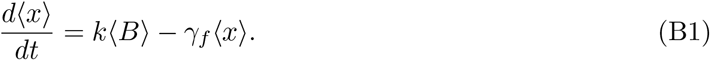

For this case, it is important to note that the dynamics of total TFs are independent of decoy sites and binding affinity. This implies that the dynamics of total TF is a result of a bursty production with rate *k* and a degradation reaction with rate *γ*_*f*_. For this busty birth and simple death process, the dynamics of the second moment can be written as,

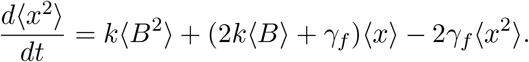

By solving (B1) and (B1) with initial condition ⟨*x*(*t*=0)⟩=0 and ⟨*x*^2^(*t*=0)⟩=0, one can obtain noise in total TF as a function of time as

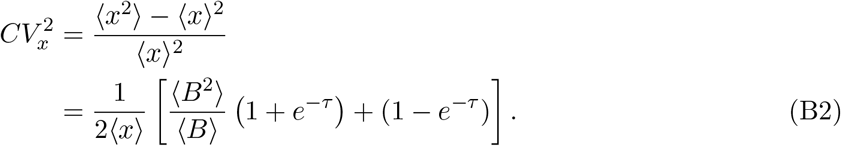

When decoy binding protects TF from decay i.e. *β*=0, dynamics of both the free TF and total TF become nonlinear, and exact calculations become challenging. However, we can obtain an approximate closed-form solution at a particular limit when the binding is very strong. In the limit of *k*_*d*_ ≪ *x*_*f*_, i.e., when all decoy sites are bound with TFs (*x*_*b*_ ≈ *N*), the dynamics of the bound TFs can be ignored. The dynamics of the first and second moments for the total TF level hence are, respectively given as following

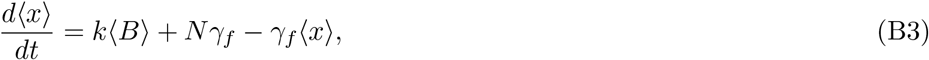

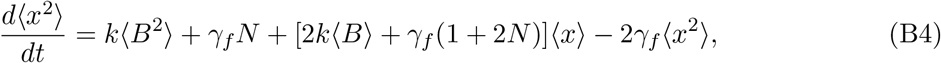

and one can exactly solve these equations with initial condition ⟨*x*^2^(0)⟩ = 0 and obtain

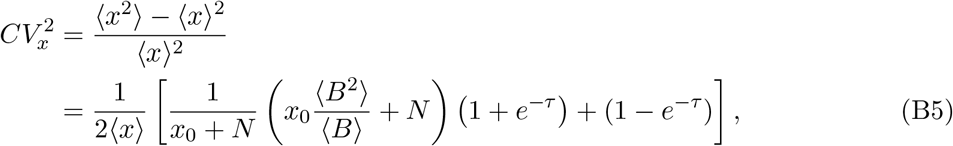

which is Eq. (21) in the main text with *τ* = *tγ*_*f*_.

**FIG. S1.**
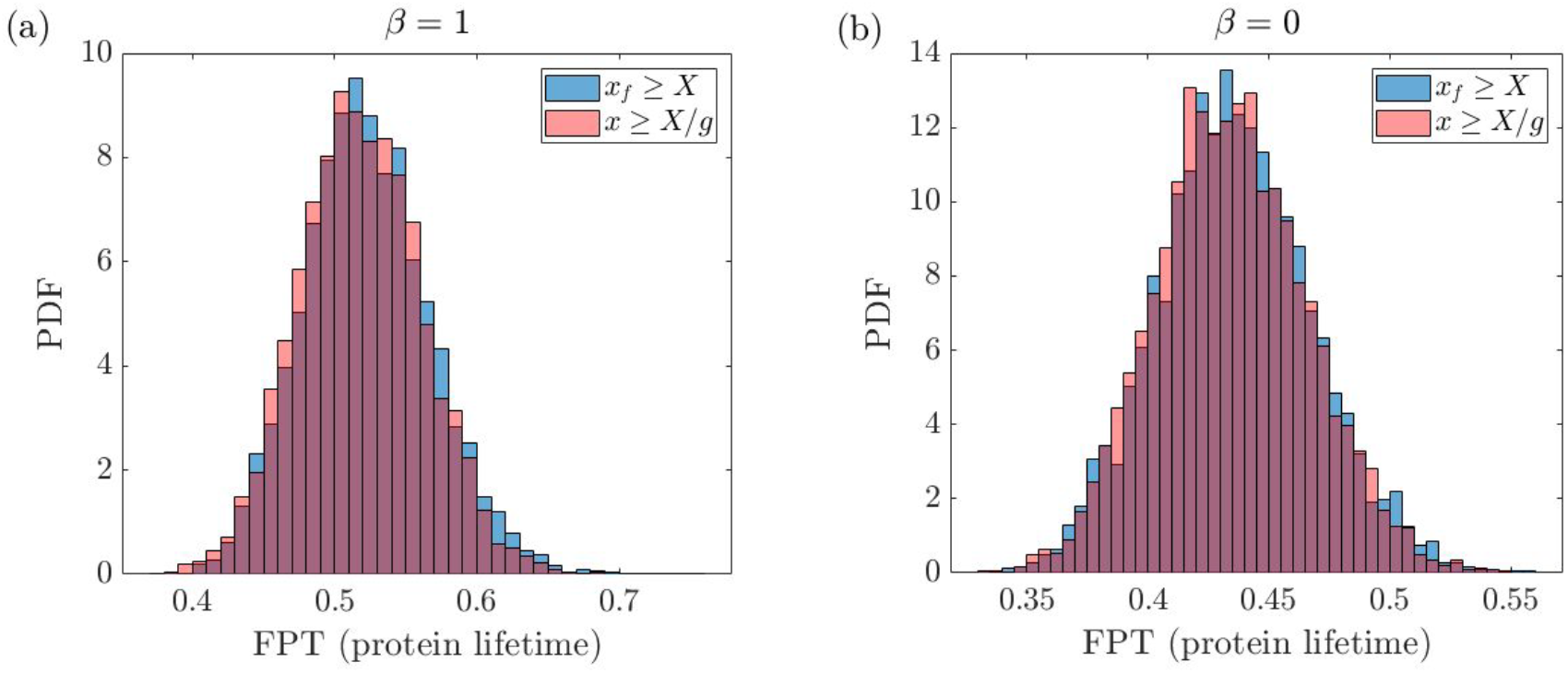
Modified definition of FPT holds even in large noise and slow binding/unbinding limit. Simulated FPT distributions from both thresholds in Eqs. (2) and (14) will show a significant overlap for **(a)** *β* = 1 and **(b)** *β* = 0. We assume the shifted geometric distribution of burst size, i.e., ⟨*B*^2^⟩ = 2⟨*B*⟩^2^ + ⟨*B*⟩ with parameter values: *γ*_*f*_ = 1, *k*_*b*_ = *k*_*u*_ = 1, ⟨*B*⟩ = 100, *x*_0_ = 1500, *N* = 300, *X* = 300. For simulation, we use the SSA (64) over 5 × 10^3^ realizations on model Eqs. (1).

**FIG. S2.**
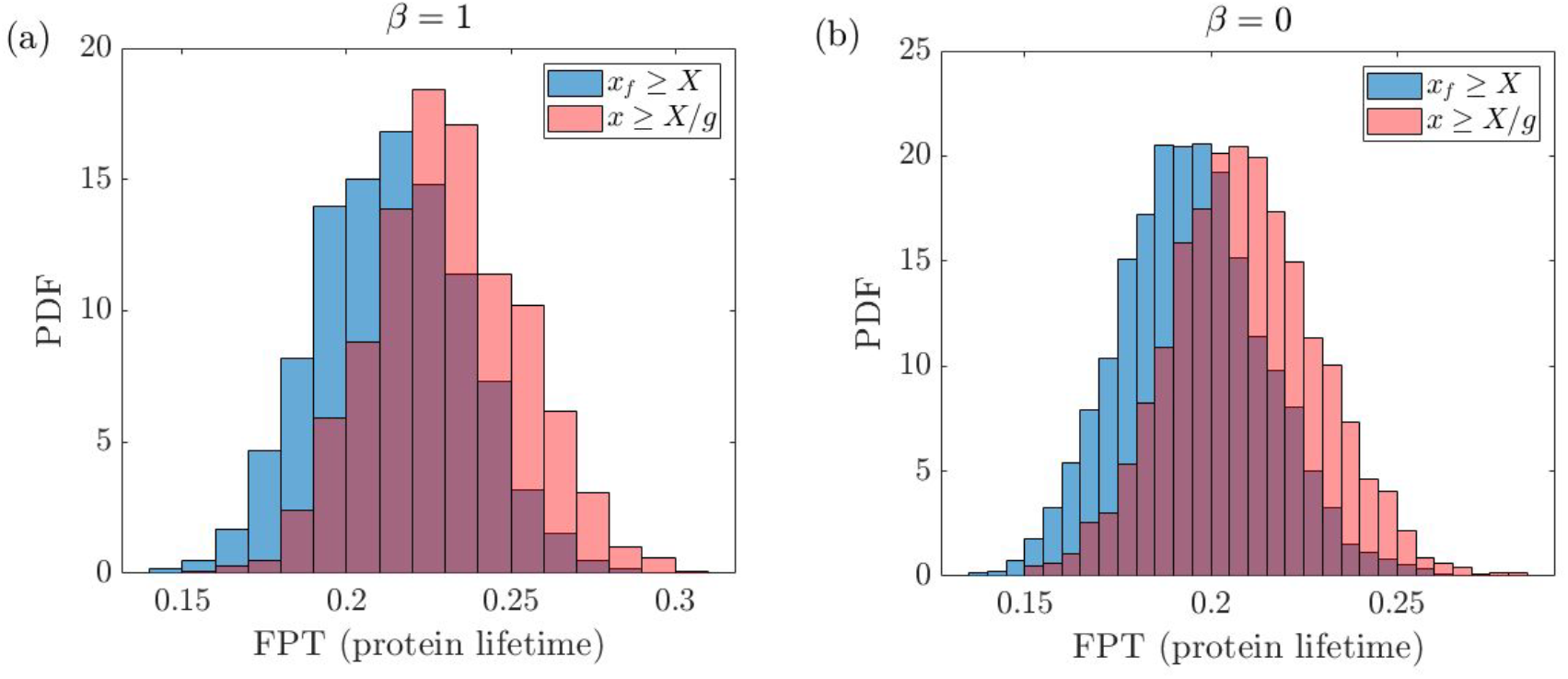
Modified definition of FPT does not hold for small threshold limit. Simulated FPT distributions from both thresholds in Eqs. (2) and (14) will show a mismatch for **(a)** *β* = 1 and **(b)** *β* = 0. We assume the shifted geometric distribution of burst size, i.e., ⟨*B*^2^⟩ = 2⟨*B*⟩^2^ + ⟨*B*⟩ with parameter values: *γ*_*f*_ = 1, *k*_*b*_ = *k*_*u*_ = 0.1, ⟨*B*⟩ = 10, *x*_0_ = 1500, *N* = 300, *X* = 15. For simulation, we use the SSA (64) over 5 × 10^3^ realizations on model Eqs. (1).

**FIG. S3.**
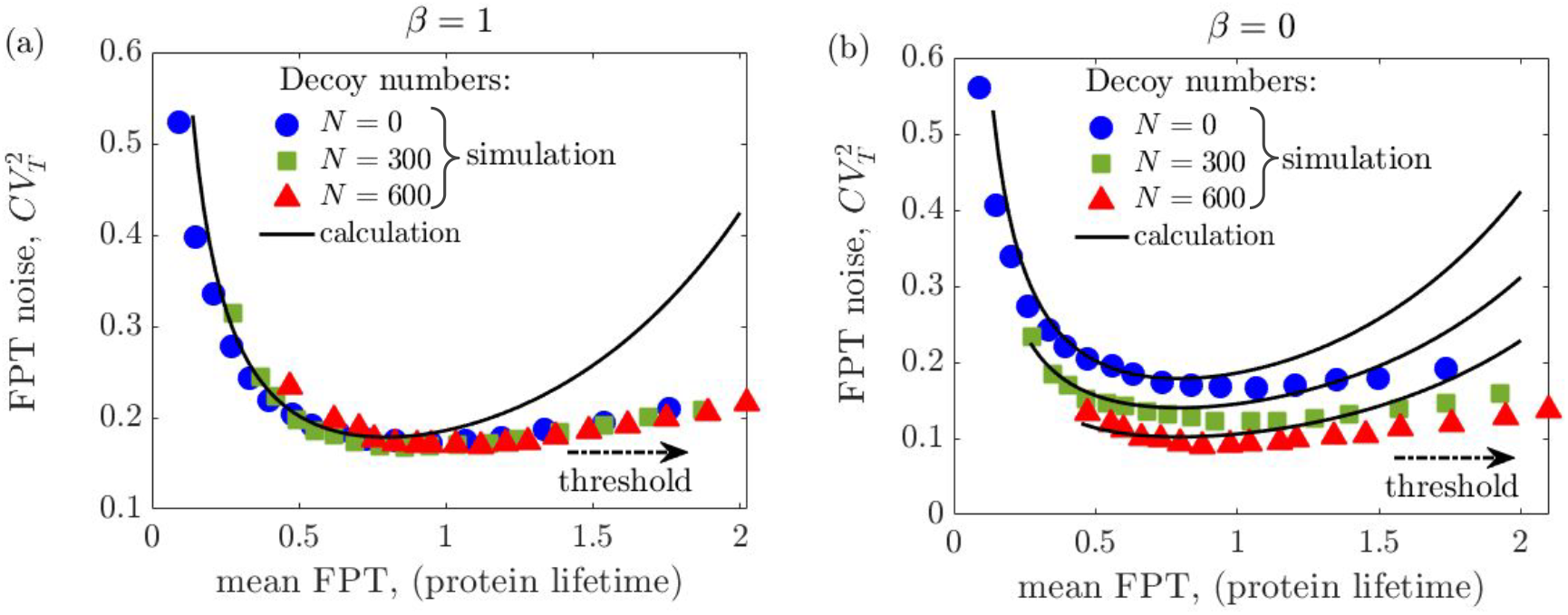
Calculation deviates from simulation in large noise and slow binding/unbinding limit. Variation of FPT noise with mean FPT for **(b)** *β* = 1 and **(c)** *β* = 0 for three different values of *N*. For *β* = 1, the relation between ⟨*T* ⟩ and 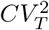 follows a master curve Eq. 18 for all *N* values, which deviates for *β* = 0, for nonzero *N* values (Eq. 22). We assume the shifted geometric distribution of burst size, i.e., ⟨*B*^2^⟩ = 2⟨*B*⟩^2^ + ⟨*B*⟩ with parameter values: *γ*_*f*_ = 1, *k*_*b*_ = *k*_*u*_ = 1, ⟨*B*⟩ = 100, *x*_0_ = 1500. For simulation, we use the SSA (64) over 5 × 10^3^ realizations on model Eqs. (1).

